# Dissecting the temporal phenomics and genomics of maize canopy cover using UAV mediated image capture

**DOI:** 10.1101/2024.06.25.600603

**Authors:** Julian Cooper, Dorothy D. Sweet, Sara B. Tirado, Nathan M. Springer, Candice N. Hirsch, Cory D. Hirsch

## Abstract

Canopy cover is an important agronomic trait influencing photosynthesis, weed suppression, biomass accumulation, and yield. Conventional methods to quantify canopy cover are time and labor-intensive. As such, little is known about how canopy cover develops over time, the stability of canopy cover across environments, or the genetic architecture of canopy cover. We used unoccupied aerial vehicle-mediated image capture to quantify plot-level canopy coverage in maize throughout the growing season. Images of 501 diverse inbred lines were acquired between 300 and 1300 growing degree days in the 2018-2021 growing seasons. We observed that the maize canopy developed following a logistic curve. Phenotypic variation in percent canopy coverage and canopy growth rate was explained by genetic and environmental factors and genotype-by-environment interactions, however the percent of variance explained by each factor varied throughout the growing season. Environmental factors explained the largest portion of trait variance during the adult vegetative growth stage and had a larger impact on canopy growth rates than percent canopy coverage. We conducted multiple genome wide association studies and found that canopy cover is a complex, polygenic trait with a diverse range of marker trait associations throughout development. The change in associations indicated that single time point phenotyping was insufficient to capture the full phenomic and genetic diversity of canopy cover in maize.

## Introduction

Canopy cover is the proportion of ground area covered by the vertical projections of above-ground plants (Jennings, 1999). In agricultural settings, the leaf area and leaf angle of surrounding plants comprise the closure over a specific plot or field (de Wit, 1965). Canopy cover is an important agronomic trait, contributing to photosynthesis, weed suppression, biomass accumulation, and yield (Jannink et al., 2001; Campillo et al., 2008; Xavier et al., 2017; García-Martínez et al., 2020). More canopy coverage early in the growing season maximizes light interception prior to the reproductive stage, allowing additional capacity for photosynthesis given favorable environmental conditions (Gifford et al., 1984; Richards, 2000). Plants with increased early season canopy coverage also capture a greater proportion of incoming solar radiation, which denies light resources to neighboring weeds during critical growth periods (Jannink et al., 2001). The benefits drawn from increased light interception has led to the selection of varieties capable of maximizing solar absorption (Board, J. E. et al., 1992; Richards, 2000).

Methods used to quantify canopy cover or light interception have historically relied on objective, time-consuming, and logistically difficult methods (Purcell, 2000; Campillo et al., 2008). Photosynthetically active radiation sensors capable of detecting visible radiation between 400 and 700 nm have been placed in fields to measure light interception by the above canopy (Board, J. E. et al., 1992). However, this method can be costly, dependent on the number of sensors needed, and is unsuitable for low-lying vegetation that must be moved or disturbed to position the sensors (Campillo et al., 2008). Visual estimates of light interception, either through quantifying vegetation cover or shadows, is a lower cost option but is prone to human error and bias (Olmstead et al., 2004). Furthermore, data must be collected during a limited timeframe surrounding solar noon with no clouds or other obstructions to minimize variance in the angle and intensity of the sun between data collection events (Purcell, 2000; Campillo et al., 2008).

New high-throughput phenotyping technologies provide researchers with novel ways to study difficult to measure phenotypes. In particular, unoccupied aerial vehicles (UAVs) have been widely adopted in recent years for their broad applicability and low barrier to entrance. UAV-mediated image capture can facilitate rapid canopy cover quantification without reliance on subjective, time-consuming measurements or specific solar and weather conditions (Sweet et al., 2022). Canopy cover has been quantified from aerial images in a variety of crops including maize, soybean, and tomatoes (Purcell, 2000; Campillo et al., 2008; Xavier et al., 2017; Makanza et al., 2018; Moreira et al., 2019; García-Martínez et al., 2020). Various image analysis techniques have been employed for measuring canopy cover. Some studies have used the Excess Green Index to identify plant pixels, however shadows and volunteer plants can confound image classification (Moreira et al., 2019; Rodene et al., 2022). Supervised machine learning algorithms can be used to increase classification accuracy under confounding conditions, however, these require manual annotation of images for classification learning (Xavier et al., 2017; Makanza et al., 2018). High-throughput phenotyping platforms and efficient image analysis pipelines also facilitate data collection at multiple time points throughout the growing season. Temporal maize canopy cover data collected by UAVs can monitor leaf senescence patterns (Makanza et al., 2018). In soybean, aerial imaging to quantify canopy cover demonstrated that early season measurements were useful for adjusting yield predictions, indicating canopy cover may be a useful secondary breeding trait for yield (Moreira et al., 2019).

Temporal data can also be incorporated into stability analyses. Static stability captures the capacity of a genotype to maintain trait performance between measured environments (Heinrich et al., 1983; Becker and Leon, 1988). Static stability can be quantified using various metrics from an additive main-effects and multiplicative interaction (AMMI) model, such as the sum across environments of absolute value of genotype by environment (GxE) interaction (AVAMGE) (Gauch, 1992; Gauch, 2013). Genotypes with small AVAMGE values have low magnitudes of GxE and a conserved phenotype across measured environments.

Conversely, dynamic stability quantifies the ability of a genotype to improve trait response as environmental conditions improve (Finlay and Wilkinson, 1963; Bernardo, 2020). Dynamic stability can be calculated using a Finlay-Wilkinson (FW) slope coefficient to measure the direction of GxE response using joint regression (Finlay and Wilkinson, 1963; Lian and de Los Campos, 2015). When modeling response to different environments, genotypes with a FW coefficient of 1.0 have the mean response to an observed environment as the population. In such cases, trait variation is likely due to environmental conditions. In contrast, slopes that are greater or less than 1.0 indicate favorable or unfavorable GxE interactions, respectively. Static and dynamic stability methods are commonly used to evaluate yield across environments, however these metrics have also been used to quantify the phenotypic plasticity of morphological traits, such as plant height and disease resistance (Fuentes et al., 2005; Mu et al., 2022). Calculating stability metrics across time can reveal how plasticity of a trait changes throughout growth and development and reveals additional information about factors affecting phenotypic variance.

When genotype is found to explain a large portion of phenotypic variance, genome wide association studies (GWAS) allow researchers to identify regions of the genome associated with traits of interest. Associated genomic regions for core maize traits, such as plant height and leaf architecture, have been identified (Buckler et al., 2009; Tian et al., 2011; Peiffer et al., 2014). These studies were fundamental to understanding maize quantitative traits, however, they mostly relied on trait measurements from a single time and did not capture temporal phenotypic diversity to test how associated genomic regions vary throughout development. Recent GWAS studies have incorporated time-resolved phenotyping, such as repeated measurements of plant height and biovolume throughout the growing season, and showed that putative causal genomic regions can vary throughout development (Muraya et al., 2017; Xavier et al., 2017; Wu et al., 2019; Knoch et al., 2020; Rodene et al., 2022). Temporal GWAS was conducted in soybean on canopy cover measurements at regular intervals from two to eight weeks after planting. By performing repeated GWAS at each time point, six genomic regions associated with canopy cover were located, including one on chromosome 19 that was significant across all days (Xavier et al., 2017).

Despite the importance of canopy cover to photosynthesis, weed suppression, biomass accumulation, and yield, there is limited research on how sources of phenotypic variation and the genetic architecture of maize canopy cover changes throughout growth and development. In this study, a UAV equipped with a red, green, and blue (RGB) camera was used to capture time-series images of a subset of the maize Wisconsin Diversity Panel (Hansey et al., 2011) throughout the 2018-2021 growing seasons. A k-means clustering algorithm and a local polynomial regression model were employed to quantify canopy cover percentage and growth rates at 50 growing degree day (GDD) intervals from 300-1300 GDD. Phenotypic variance partitioning and static and dynamic stability estimates were calculated for canopy cover traits throughout the growing season. Finally, a genome wide association study was conducted using phenotypes from multiple modeling approaches for each trait-time iteration.

## Results and Discussion

### An unsupervised k-means clustering pipeline to measure canopy cover

To quantify in field maize canopy cover over time, a UAV equipped with an RGB camera was used to image 501 inbred lines from the Wisconsin diversity panel (Hansey et al., 2011) throughout the 2018, 2019, 2020, and 2021 growing seasons. K-means was used to cluster pixels and identify plants from the images (Figure 1). K-means clustering is an unsupervised algorithm that facilitated flexible and effective image masking. To accurately extract canopy cover at multiple time points, the number of clusters was manually adjusted to account for variable flight conditions. For example, partially cloudy skies caused variation in the brightness and hue saturation between sunny and shaded areas within the field. If flights were conducted at early or late times of day the sun conditions could produce long shadows from taller plants that would be cast on nearby plots and cause variation in the color appearance of plant pixels. By altering the number of clusters used for image classification in response to flight conditions, plant pixels were successfully masked and non-plant pixels were removed without having to create training data for masking.

**Figure 1.**
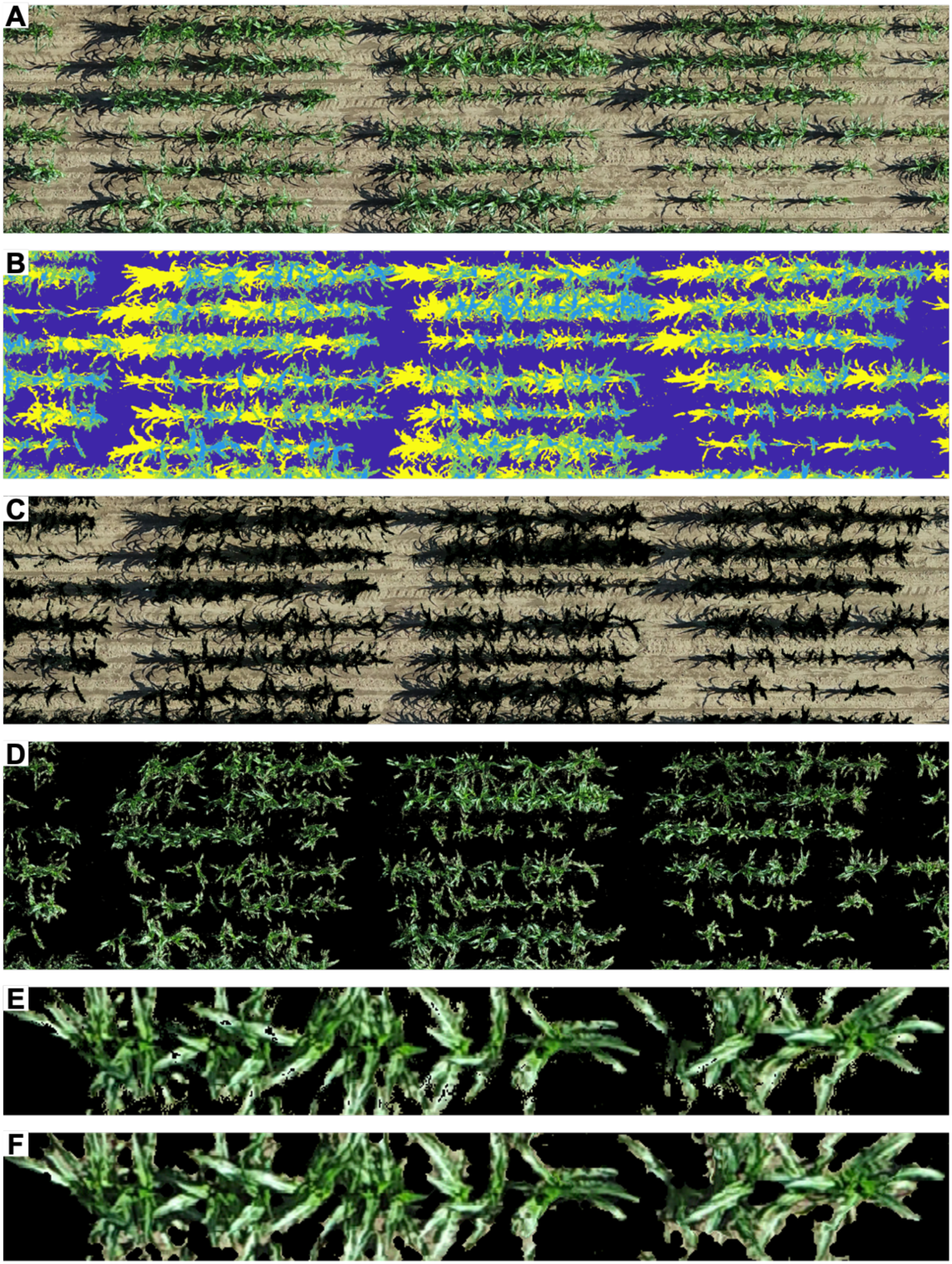
K-means clustering method used to extract canopy cover from UAV captured images. A. Subsection of a field-level reconstructed orthomosaic. B. Pseudo colored image of the field subsection showing four k-means clusters based on Red Green Blue (RGB) pixel values. C. K-means clusters containing non-plant associated pixels are shown in RGB colors. These pixels were excluded from further analysis. D. K-means clusters containing plant-associated pixels are shown in RGB colors. These pixels were used in downstream analysis. E. Plot level example of a masked image, prior to image dilation showing holes in vegetation masking. F. Plot level example of dilated image that produced the final mask used to quantify canopy cover as the percentage of masked plant pixels out of the total number of plot pixels.

A limitation of using k-means clustering was the inability to differentiate types of plant material. When non-maize plants were present in images, green pixels of maize, weeds, or volunteer crops were often included in the same cluster, which would inflate maize canopy cover values. Making distinctions between plant types is a difficult problem for plant imaging pipelines and is also an issue when using other masking pipelines, including using the Excess Green Index (Meyer and Neto, 2008; Rodene et al., 2022). To ensure the accuracy of canopy cover estimations, plots with weed pressure were removed from subsequent analysis. The inclusion of plant height into future canopy cover quantification pipelines may be a solution for excluding low-lying weeds in plots to minimize data loss.

### Maize canopy cover development followed a logistic growth curve

Using the k-means clustering pipeline, canopy cover was extracted from 300 to 1300 GDD. A locally estimated scatterplot smoothing (LOESS) polynomial regression was used to predict canopy cover for each plot at 50 GDD intervals to allow comparisons at equivalent growth stages between environments (Supplemental Figure S1, Supplemental Table S1). Local regression methods fit simple models to localized subsets of data to preserve variation that would be lost using other regression techniques. This flexibility makes LOESS ideal for fitting biological canopy cover data while preserving differences across time and between environments (Van Tassel et al., 2022). Local polynomial regression relies on a minimum threshold of observations and can be imprecise when there are large intervals between observations or extreme outliers. Using regularly spaced UAV flights for time-series phenotyping and effective data quality control mitigated these drawbacks in our phenotyping pipeline.

Maize canopy coverage development followed a logistic growth curve (Figure 2A). This pattern has been observed in maize and soybean canopy coverage and maize height developmental curves (Xavier et al., 2017; Makanza et al., 2018; Malambo et al., 2018; Tirado et al., 2020). The phases of the fitted logistic curve corresponded to important maize developmental stages. Maize canopy coverage increased slowly from 300 GDD to 450 GDD (Supplemental Table S2, Figure 2A), which corresponds to the estimated timing of juvenile vegetative growth and marks the transition from seed energy stores to the beginning of photosynthesis derived energy (Abendroth et al., 2011). At approximately 500 GDD, the plants entered a period of rapid canopy growth and canopy cover increased at an average rate of 0.08% per GDD, which equates to about 1.8% per calendar day (Supplemental Table S2, Figure 2A). This timing is consistent with the expected onset of maize rapid growth during the adult vegetative stage of development, where new leaf stages can occur every 1-2 days (Abendroth et al., 2011; Malambo et al., 2018). Maize canopy coverage began to stabilize around 1100 GDD, which marks the beginning of the reproductive phase when tassels emerge and leaf production terminates (Abendroth et al., 2011) (Supplemental Table S2, Figure 2A). In the Wisconsin Diversity panel, the average flowering time was 1300 GDD (Hansey et al., 2011) and to avoid phenotypic variance due to senescence, 1300 GDD was chosen as the terminal point for canopy cover for all years.

**Figure 2.**
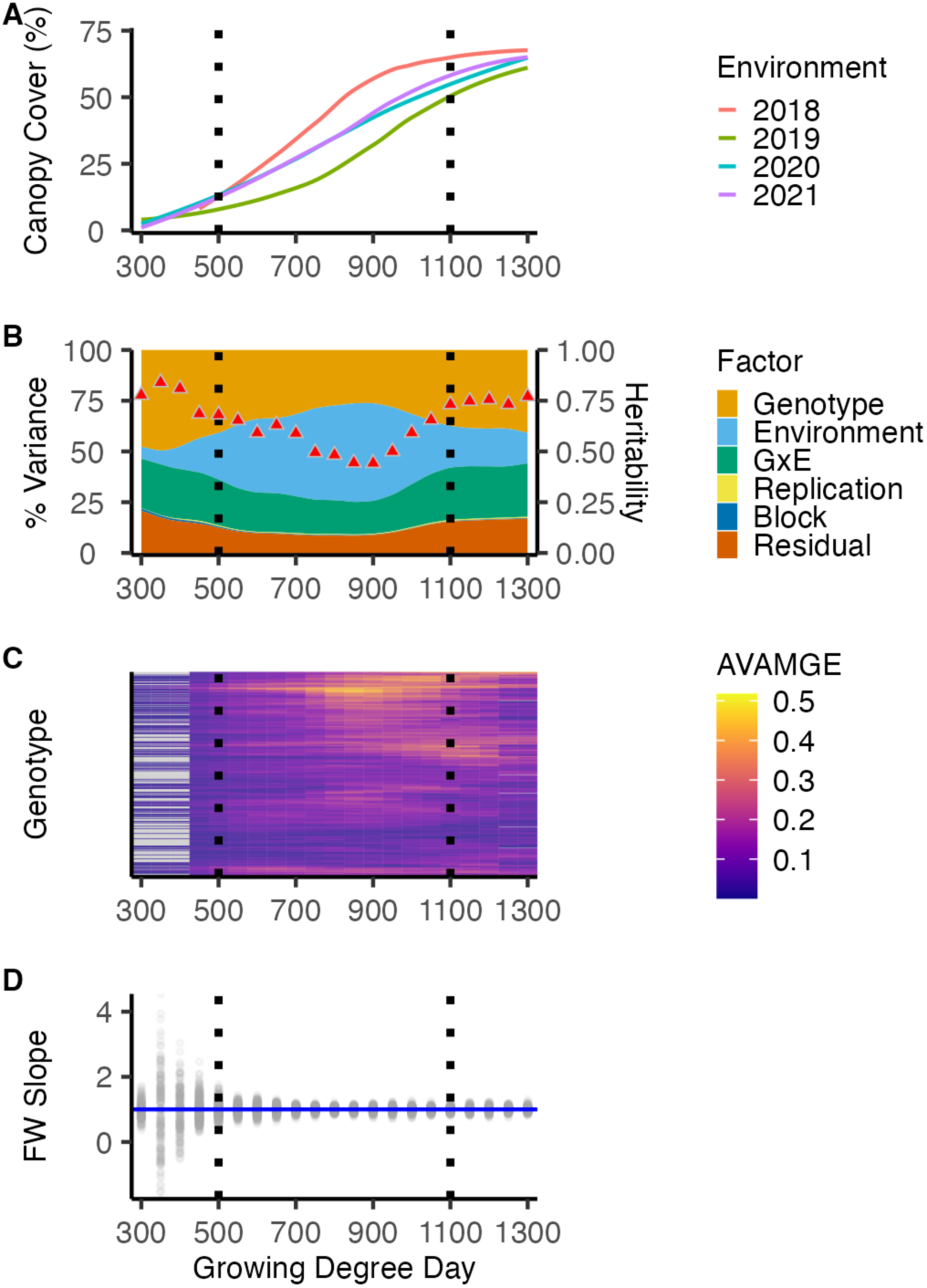
Temporal canopy cover traits and variance partitioning. Juvenile vegetative, adult vegetative, and reproductive developmental stages are partitioned by dashed black lines from left to right in each of the plots, respectively. A. The average canopy growth curve for each environment based on local regression predicted time points. B. The percentage of variance explained (PVE) by different experimental factors for percent canopy coverage. Heritability on a line-mean basis from 0.0-1.0 is shown as a red triangle at each time point. C. The sum across environments of absolute value of genotype by environment interaction (AVAMGE) modeled by an additive main-effects and multiplicative interaction (AMMI) model for percent canopy coverage. Genotypes with less than the minimum of five replicates across three environments are designated as missing data with gray color. D. The range of Finlay Wilkinson (FW) joint regression coefficients for percent canopy coverage. The blue line marks a FW slope of 1.0.

### Timing of factors that influenced canopy cover variation

Statistically significant differences were observed between mean terminal canopy coverage between three of the four environments (Table 1). However, the coefficients of variation (CV) used to scale canopy coverage variance across GDD were higher during juvenile and adult vegetative growth than in the reproductive phase (Supplemental Table S2). Therefore, more phenotypic variation was captured from in-season canopy coverage measurements than as a terminal trait. To determine how the factors that influenced canopy cover variation changed over time, an analysis of variance (ANOVA) was conducted to calculate the percent of variance explained (PVE) by genotype, environment, GxE, replication, block, and experimental error, along with the broad-sense heritability at each GDD (Figure 2B, Supplemental Table S2). All of the factors tested were statistically significant (p < .05) except replication at 300 and 350 GDD and block at 1000 GDD. The PVE from block or replication never exceeded 1.2%, which indicated the observed differences between canopy cover were not due to confounding experimental error such as spatial variation between replications or technical error from the UAV or image analysis pipelines. At terminal canopy coverage, the PVE by genotype, environment, and GxE was 41.9%, 13.9%, and 26.8%, respectively, however the influence of these factors varied throughout development.

**Table 1.** Yearly weather from 300-1300 Growing Degree Days (GDD) and mean terminal canopy coverage for each environment. Terminal canopy coverage is at 1300 GDD, the average flowering time for the Wisconsin Diversity Panel. A Tukey Honestly Significant Difference (HSD) test was performed to determine if the mean canopy cover percentage differed between environments. Tukey HSD test groups identify environments with no statistical difference between mean terminal canopy coverage.

| Year | Planting Date | Days to 1300 GDD | Average Minimum Temp (F) | Average Maximum Temp (F) | Total Precipitation (in.) | Average Terminal Canopy Cover (%) | Tukey HSD Test Group |
| --- | --- | --- | --- | --- | --- | --- | --- |
| 2018 | 5/14/18 | 58 | 67.4 | 83.1 | 9.59 | 69.0 | A |
| 2019 | 5/30/19 | 65 | 62.1 | 79.5 | 12.2 | 60.8 | B |
| 2020 | 5/7/20 | 75 | 61.6 | 81.3 | 5.42 | 64.5 | C |
| 2021 | 5/6/21 | 75 | 63.3 | 84.4 | 3.01 | 64.9 | C |

#### Genotype explained the most phenotypic variance during juvenile vegetative and reproductive growth stages

During juvenile vegetative growth and the reproductive phase, genotype explained the highest proportion of observed phenotypic variance (Figure 2B). Maize seedling vigor and early season leaf architecture are known to have a large genetic component, which could explain the high PVE by genotype early in the season (Li et al., 2022; Wang et al., 2022). For maize plant height, a related continuous trait, the PVE by pedigree also varied throughout the growing season and was largest during the reproductive growth stage (Adak et al., 2021). The heritability of canopy cover percentage was highest when the PVE by genotype was large (Figure 2B). The high heritability and large PVE by genotype during these times means breeders may be successful in directly selecting for canopy coverage as trait, whether to increase early season canopy coverage to maximize weed suppression or to decrease terminal coverage to increase planting densities and potentially yield.

#### The environment had the most impact on phenotypic variance during adult vegetative growth

It has been shown that growing conditions during maize adult vegetative growth can impact plant height and leaf area (Dodig et al., 2021). Throughout adult vegetative growth, the environment contributed the most PVE for maize canopy cover and the heritability was lower than during the juvenile vegetative or reproductive stage (Figure 2B). Biologically, this appears to indicate that variation in maize canopy cover was more due to phenotypic plasticity than genotypic differences during the adult vegetative growth stage. Some portion of canopy variation was likely due to independent weather patterns between environments, such as differences in mean temperatures and total precipitation (Table 1). Some phenotypic variation may also be due to temporal variation in weather conditions (i.e. timing and magnitude of different precipitation events) (Supplemental Figure S2A). For example, 2018 had the highest average minimum temperature and most precipitation events during the adult vegetative growth stage, and plots achieved peak canopy coverage earlier and had the highest mean terminal canopy coverage of all environments (Table 1, Supplemental Figure S2B). Similarly, in sorghum the diurnal temperature range during the period of rapid growth from 40-53 days after planting was highly influential on plant height (Mu et al., 2022). Due to the high PVE by environment during the adult vegetative growth stage, physiologists seeking to model maize canopy coverage may benefit from using environmental indices generated during that time to improve predictions. Furthermore, because these environmental indices can be calculated in-season, they may be useful for predicting end of season performance in untested environments (Li et al., 2018).

#### The magnitude and direction of GxE interactions varied between inbreds across development

In addition to genotype and the environment, we observed a significant PVE by GxE interactions throughout maize development, ranging from 13.7-26.8%. Different magnitudes and directions of GxE can be exploited to maintain or maximize performance across environments. To investigate how the magnitude of GxE interactions varied between genotypes and across time, AVAMGE values from an AMMI model were used to quantify static stability. In the adult reproductive phase, certain genotypes had increased AVAMGE values relative to the rest of the population, which indicates high phenotypic variation between environments (Figure 2C, Supplemental Table S2). AVAMGE are observation specific, so direct comparisons of the magnitude of GxE between GDD are not possible using an AMMI model, however the high CV throughout growth stages indicates that a wide range of GxE interactions were captured for different genotypes throughout development.

The differences in static stability between genotypes is evident when comparing growth curves between environments. For example, inbred I29 had the highest mean AVAMGE values across all GDD, and canopy growth curves were variable across time and between years, particularly in 2018 (Supplemental Figure S3A). Between plots of I29 in different environments at a given GDD, there was a maximum difference in canopy cover of 50% (Supplemental Figure S3A). In contrast, inbred PHM10 had the lowest average AVAMGE and demonstrated conserved growth patterns between environments, with a maximum difference between plots of 36% (Supplemental Figure S3B). Notably, the range of terminal canopy cover values was similar for both genotypes, despite differences in static stability. The majority of differentiation between genotypes was observed in the adult vegetative growth phase. Thus, differences in the static stability of these two inbreds may have been missed if canopy performance was evaluated using only terminal measurements. For breeders seeking to ignore or reduce GxE and identify top performing germplasm across all desired environments, temporality adds a new dimension to data to improve selection of genotypes with stable performance across time, not just at the end of growing season.

In addition to static stability, GxE interactions can be measured using FW slope coefficients to estimate the dynamic stability of genotype, or how performance relates to growing conditions. The range of FW values was largest during juvenile vegetative growth, with a range of favorable and unfavorable responses to different environments as evidenced by the number of slope coefficients appreciably greater or less than 1.0 (Figure 2D). Throughout the adult vegetative and reproductive stages, the range of FW slopes constricted around 1.0, indicating that genotypic response to each environment was similar to the mean population response (Figure 2D). This corroborates the high PVE by environment seen during the adult vegetative growth stage (Figure 2B). The range of FW slope values during the adult vegetative and reproductive growth stages were similar to maize plant height (Falcon et al., 2020), although this study only used 31 inbred lines compared to our larger diversity panel.

The absolute values of FW slope coefficients of each genotype were averaged across time to compare growth curves between inbreds with low and high dynamic stability. Inbred YE 4 had the lowest mean FW slope (Supplemental Figure S3C). Plots of this genotype performed similarly in all environments, regardless of mean population canopy cover in each environment (Table 1). Inbred N192 had the highest mean FW slope across time (Supplemental Figure S4D). This genotype had conserved growth curves within environments, but different patterns between environments. As expected based on dynamic stability estimates, N192 performed the best in 2018, which had the highest mean canopy cover across all genotypes (Table 1). However, when comparing terminal canopy cover values for N192, the range of final percentages was just 22%.

What truly differentiated N192 was that plots in 2018 achieved maximum canopy coverage earlier than the other environments. This is most emphasized when comparing growth curves between 2018 and 2019 (Supplemental Figure S3D). Thus, even though plots achieved similar terminal canopy coverage, the high dynamic stability of N192 allowed this genotype to take advantage of favorable growing conditions in 2018 and maximize early season growth, which can have benefits for weed suppression and increased light interception. For breeders who wish to exploit GxE and identify elite germplasm for specific environments, growth curves can provide an additional metric to refine selection and take advantage of favorable in-season differences in growing conditions.

To determine if maize canopy accumulation was related to stability, inter-year best linear unbiased predictions (BLUPs) were generated for each genotype and compared to FW slope and AVAMGE values (Supplemental Figure S4A). The pairwise correlation between inter-year BLUPs and FW joint regression coefficients was high during the adult vegetative growth phase, which shows that canopy accumulation was related to the individual’s ability to optimize growth under improved environmental conditions. Otherwise the correlation between the three modeling approaches was low. This is expected for static and dynamic stability, as the ability to perform consistently across environments is fundamentally opposed to the ability to improve performance in optimal growing conditions (Becker, 1981). Previous studies exploring quality traits in winter barley and yield in common beans also observed a low degree of relatedness between static stability and mean performance (Kefelegn et al., 2016; Knapp et al., 2017). This suggests that it may be possible for breeders to select for elite cultivars capable of maximizing stability and performance. In summary, the impact of GxE interactions on maize canopy development was highly variable between inbreds and throughout growth and development. We hypothesize that maize canopy cover has an intrinsic floor and ceiling, defined by individual genetics, however development along that spectrum is influenced by environmental conditions and unique GxE interactions, particularly during adult vegetative growth.

### The rate of canopy growth was more affected by the environment than percent canopy coverage

Growth rates can be used to further understand phenotypic variation and responsiveness to different environments for continuous traits, such as maize plant height (Tirado et al., 2020). The rate of canopy development was quantified by calculating the slope between each LOESS predicted canopy cover percentage (Figure 3A). To test the relatedness between canopy percent and rate, the correlation at each time point was calculated (Supplemental Figure S5). In addition, PVE, heritability, AVAMGE, and FW values were calculated to determine if factors that influenced the rate of canopy development were similar in magnitude and temporality to those impacting percent canopy coverage (Figure 3B, 3C, 3D). For both traits, genotype had the largest PVE during juvenile vegetative growth, and the correlation between percent and rate was highest during this stage (Figure 2B, Figure 3B, Supplemental Figure S5). This supports the hypothesis that initial maize canopy coverage is heavily influenced by genotype. The PVE by environment was greater and heritability was less for rate of canopy development than percent canopy cover in all growth stages (Figure 3B). The correlation between percent and rate declined during the adult vegetative phase, when the environment had the most PVE (Figure 2B, Figure 3B, Supplemental Figure S5). The high importance of the environment for both canopy traits suggested that growing conditions play an important role in canopy trait variation. However, the low correlations suggested that environmental factors such as temperature and precipitation may impact the rate of canopy growth and percent canopy cover differently.

**Figure 3.**
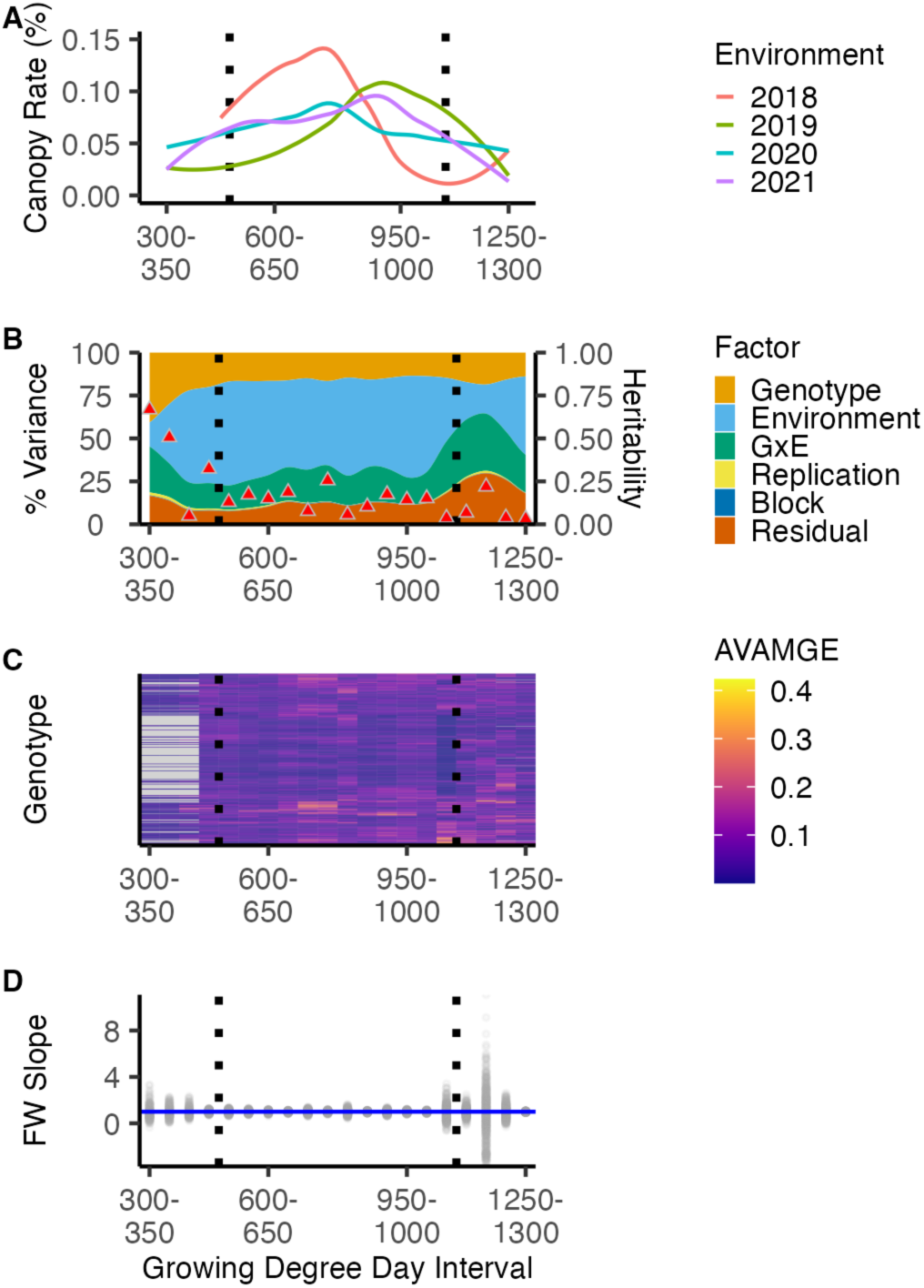
Rate and variance partitioning of canopy development of every 50 growing degree day (GDD) interval. Juvenile vegetative, adult vegetative, and reproductive growth stages are separated by dashed black lines from left to right, respectively. A. The distribution of the rate of canopy development per plot for all growing seasons between each local regression predicted time point every 50 GDD. Colored lines represent the average rate of canopy growth in each environment. B. The percent of variance explained (PVE) by factor for the rate of canopy development. The heritability on a line-mean basis from 0.0-1.0 is shown as red triangles for each Growing Degree Day interval. C. The sum across environments of absolute value of genotype by environment interaction (AVAMGE) modeled by an additive main-effects and multiplicative interaction (AMMI) model for rate of canopy development. Genotypes with less than the minimum of five replicates across three environments are colored gray to designated missing data. D. The range of Finlay Wilkinson (FW) joint regression coefficients for the rate of canopy development. The horizontal blue line marks a FW slope of 1.0.

The large PVE by environment resulted in an increase in the diversity of GxE interactions for canopy growth rates compared to canopy cover percentage. The range of static stability, measured using the CV of AVAMGE values, was greater at all time points for rate of canopy development than percent canopy cover (Supplemental Table S2). Furthermore, certain genotypes had increased AVAMGE values during specific but disjointed GDD intervals, such as 750-850 GDD and 1050-1100 GDD (Figure 3C). These differences between GDD intervals likely indicated that temporal variation in weather between environments had a large impact on phenotypic variation. Previously it was demonstrated that measurements of static stability are highly specific to populations, traits, and environments (Knapp et al., 2017; Shojaei et al., 2021). Our results further indicated that even between traits derived from the same data, different magnitudes of GxE interactions were captured for traits highly sensitive to environmental conditions. In addition, AMMI values correlated poorly with inter-year BLUP for canopy growth rates (Supplemental Figure S4B), suggesting the magnitude of GxE was not directly related to genotype performance for canopy growth rates.

Dynamic stability estimates confirmed the impact of temporal variation in weather on GxE interactions in canopy cover. There was a wider range of FW slope coefficients during adult vegetative and reproductive growth for rate than percent canopy cover, and the correlation between inter-year BLUPs and FW slopes oscillated between highly positive, no relationship, and highly negative across time points (Supplemental Table S2, Supplemental Figure S4B). The range of favorable and unfavorable GxE interactions and the fluctuations between dynamic stability and canopy growth rate demonstrates how temporal variation in weather between years affected trait variation, even between adjacent GDDs. This could be due to genotypes being optimized for specific weather conditions, but an alternative causal mechanism for diverse GxE interactions may be predicated on the speed at which growth alters in response to new conditions.

The high PVE by environment and the subsequent increase in diversity of GxE interactions resulted in higher phenotypic variation in rate of canopy cover within and across GDD intervals than was observed for percent canopy cover. Within GDD intervals, the CV for the rate of canopy development was higher than the CV for percent canopy coverage in all growth stages (Supplemental Table S2). Across GDD intervals, we expected the rate of canopy development to follow a bell curve across time based on the conserved canopy cover logistic growth curve. This was not the case, however (Figure 3A). The rate of canopy development did not increase at a constant rate in the juvenile vegetative growth phase, peak during the adult vegetative stage, and decline through the onset of the reproductive stage until terminal canopy coverage at 1300 GDD. Instead, the range of canopy growth rates varied widely between environments and between adjacent GDD intervals in the same environment. For example, the GDD interval with the peak rate of canopy development varied between environments. In 2018 (750-800 GDD), 2020 (700-750 GDD), and 2021 (800-850 GDD), the peak rate of canopy growth was early in the adult vegetative phase, whereas in 2019 (950-1000 GDD) the peak occurred near the end of the adult vegetative growth phase (Supplemental Figure S2C). Early season temperatures were low in 2019, however an increase in the average maximum temperature and a large precipitation event that preceded 950 GDD may explain why rate of canopy development peaked later (Supplemental Figure S2A).

The increased inter- and intra-season variation of canopy growth rates compared to percentages provides a unique opportunity to study cause and effect relationships of environmental factors. For example, water, temperature, light quality, and available soil nutrients are known to impact canopy area and vegetative production (Markham and Stoltenberg, 2010; Entringer et al., 2014; Laza et al., 2015; Hatfield, 2016; Rodene et al., 2022). By timing UAV flights directly following environmental events, high temporal resolution data at the onset and short and long-term impacts on maize growth can be obtained. Unlike using terminal plant measurements, or even in-season canopy cover percentages, highly environmentally sensitive traits such as growth rates could better reveal the temporal impacts of the environment on maize canopy coverage.

### The genetic architecture of maize canopy cover

To identify putative causal regions of the genome associated with canopy cover, multiple GWAS were conducted. As the ANOVA results indicated the environment was significant (p < .05), intra-year BLUPs for canopy cover percentage and rate were calculated at each time point for use as phenotypes for GWAS. In addition, for each trait-time iteration, inter-year BLUPs, FW slope coefficients, and AVAMGE values were used as phenotypes to identify potential regions of the genome involved with phenotypic plasticity and GxE interactions. In total, 326 unique trait-model-time iterations were used.

We found that maize canopy coverage was a complex, quantitative trait influenced by many small-effect loc (Figure 4). After filtering for markers in linkage disequilibrium, 1,368 significant marker trait associations (MTA) were identified across all observations (Table 2). These MTA comprised 864 unique single nucleotide polymorphisms (SNPs), located across the 10 maize chromosomes with an average absolute value of a normalized effect size of 0.07 (Supplemental Table S3). By using different trait measurements, modeling approaches, and temporal observations, we found that many distinct regions influenced the phenotypic variation observed for canopy cover. Of the unique SNPs identified, 633 were distinct to one trait-model-time iteration while 231 SNPs were identified in multiple MTAs (Supplemental Figure S6). The SNP rs277942181 had 13 MTA associated with percent canopy cover, which was the most associations for all SNPs. This SNP is in the intron of the NHL-domain containing protein coding gene Zm00001eb253510 on chromosome 5 (Woodhouse et al., 2021) (Supplemental Table S3). This gene is classified as an integral cellular membrane component and proteins containing NHL domains. NHL-domain containing proteins in plants are versatile regulators involved in various physiological and organ developmental processes, particularly in response to environmental stresses and pathogen attacks (Bao et al., 2016). Their ability to mediate protein-protein interactions and participate in signaling pathways makes them crucial components of plant adaptive mechanisms.

**Figure 4.**
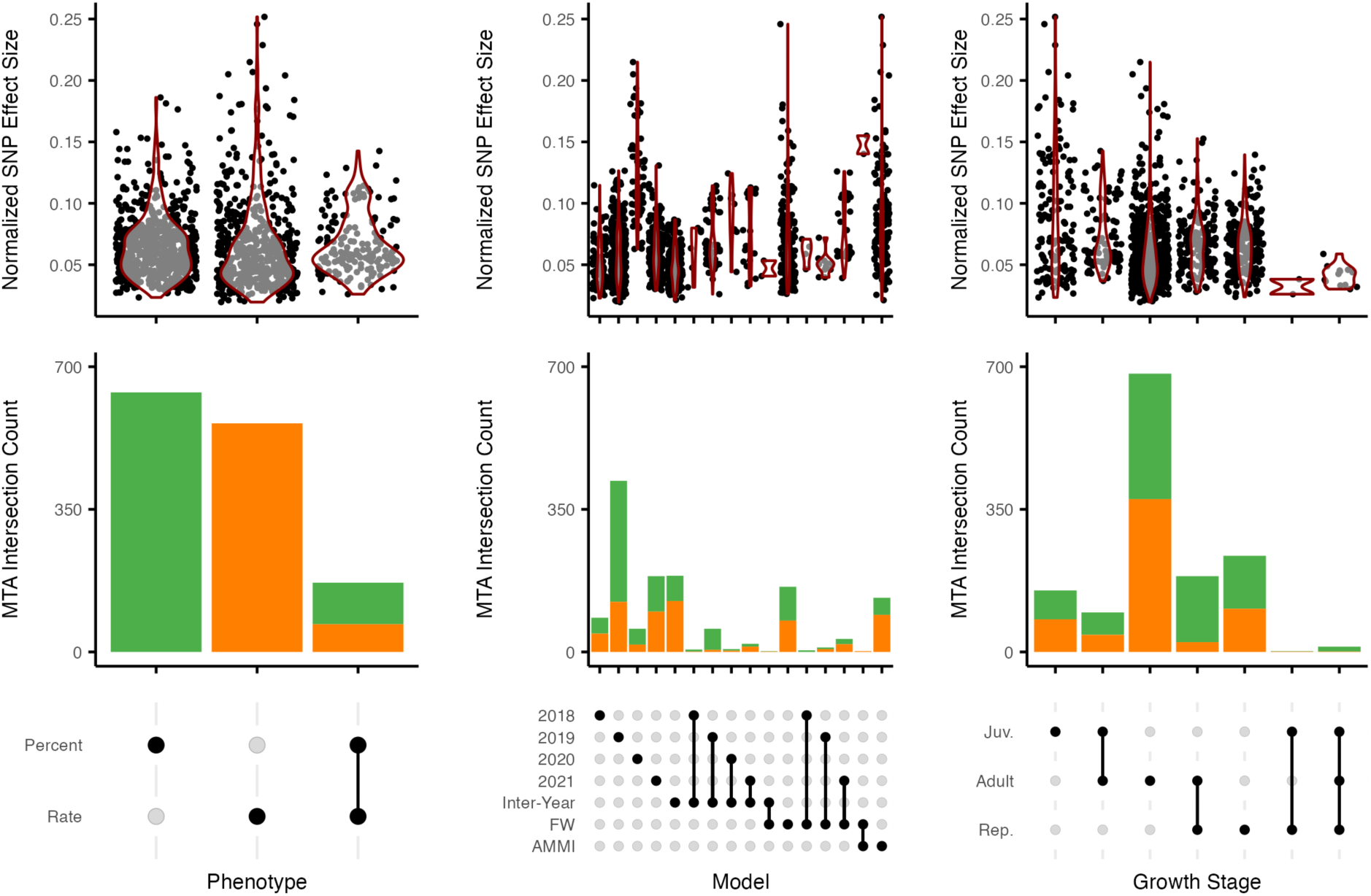
Absolute values of the normalized effect size of significant marker trait associations (MTA) and intersection of MTAs across canopy traits, modeling approaches, and growth stages. Percent canopy cover (Percent) MTA are shown in green and rate of canopy development (Rate) MTA are shown in orange. A. Absolute values of the normalized effect size from all models and time points for Percent and Rate. B. Intersection of MTA from all models and time points for Percent and Rate. C. Absolute values of the normalized effect size from all traits and time points for intra-year linear (2018, 2019, 2020, 2021), inter-year linear, Finlay-Wilkinson (FW) joint regression, and additive main-effects and multiplicative interaction (AMMI) models. D. Intersection of MTA from all traits and time points for intra-year linear (2018, 2019, 2020, 2021), inter-year linear, Finlay-Wilkinson (FW) joint regression, and additive main-effects and multiplicative interaction (AMMI) models. E. Absolute values of the normalized effect size from all traits and models for the juvenile vegetative (Juv.), adult vegetative (Adult), and reproductive (Rep.) growth stage. F. Intersection of MTA from all traits and models for the juvenile vegetative (Juv.), adult vegetative (Adult), and reproductive (Rep.) growth stage.

**Table 2:** The counts of total marker trait associations (MTA) and unique single nucleotide polymorphisms (SNP) for all observations and each phenotype [percent canopy cover (Percent), rate of canopy development (Rate)], model [intra-year linear (2018, 2019, 2020, 2021), inter-year linear, Finlay-Wilkinson (FW) joint regression, and additive main-effects and multiplicative interaction (AMMI)], and growth stage [juvenile vegetative (Juv.), adult vegetative (Adult), and reproductive (Rep.)]

| Category | Class | Total MTA | Unique SNPs |
| --- | --- | --- | --- |
| ALL | ALL | 1368 | 864 |
| Phenotype | Percent | 739 | 393 |
| Phenotype | Rate | 629 | 500 |
| Model | 2018 | 91 | 61 |
| Model | 2019 | 464 | 239 |
| Model | 2020 | 61 | 37 |
| Model | 2021 | 227 | 112 |
| Model | Inter-Year | 219 | 172 |
| Model | FW | 172 | 153 |
| Model | AMMI | 134 | 121 |
| Growth Stage | Juv. | 188 | 150 |
| Growth Stage | Adult | 865 | 558 |
| Growth Stage | Rep. | 315 | 223 |

#### Different genomic regions are associated with maize canopy coverage and rate of canopy growth

To identify shared and distinct regions of the genome between canopy traits, separate GWAS were conducted for percent canopy cover and growth rate across all models and time points. The regions of the genome responsible for controlling canopy coverage were different from loci controlling the rate of canopy growth, although the effect size of markers for both traits was similar (Figure 4A, 4B). This was expected due to the low correlation between the two traits (Supplemental Figure S5). There were 393 unique significant SNPs out of 739 total MTA found for percent canopy cover and 500 unique significant SNPs out of 629 MTA for rate of canopy development (Table 2). Only 33 unique significant SNPs out of 170 MTA were shared between percent and rate.

Distinct regions of the genome can be associated with multiple related traits. For example, plant height and canopy height in cotton had about 50% of identified significant loci shared between the two traits, although the traits had a strong correlation of 0.91 (Pauli et al., 2016). The low correlation and distinct regions of the genome for temporal canopy traits provides multiple avenues for plant breeders targeting maize canopy coverage. For example, separate favorable quantitative trait loci (QTL) could be stacked to increase both the amount and speed of biomass accumulation.

#### Different modeling approaches identified unique SNPs associated with phenotypic plasticity between environments

Historically little relationship between trait performance, static stability, and dynamic stability has been found (Becker, 1981; Kefelegn et al., 2016; Knapp et al., 2017). For example, a study comparing SNPs associated with soybean yield identified by BLUPs versus various stability models found only 1 out of 86 independent QTL shared between the modeling approaches (Happ et al., 2021). Separate QTL identified by BLUPs and stability metrics can be used to explain novel genetic and GxE variance. We used different modeling approaches to identify distinct and potentially shared regions of the genome associated with maize canopy coverage in specific environments (intra-year BLUPs), across environments (inter-year BLUPs), and stability (FW and AMMI). No SNP was detected in more than two models (Figure 4D). In particular, there was little overlap in MTA shared between inter-year BLUPs, AMMI values, or FW coefficients. This was expected, as the correlation across time between these modeling approaches was generally low for both traits (Supplemental Figure S4). Interestingly, the average effect size of inter-year BLUPs was less than the effect size of MTA for stability metrics (Figure 4C). This appears to show that averaging performance across environments to calculate BLUPs resulted in a loss of GXE variation compared to explicitly measuring conserved performance using static stability estimates or adaptive performance using dynamic stability estimates. These findings support previous theories that different regions of the genome are associated with different components of a trait of interest which provides opportunities for researchers to dissect the different causal mechanisms affecting performance versus stability (Happ et al., 2021).

Within individual environments there was variation in the number of unique SNPs and MTA identified using intra-year BLUPs (Table 2). No MTA were shared between individual environments (Figure 4D). The effect size of canopy cover MTAs was constant in 2018, 2019, and 2021, despite fewer MTA identified in 2018 and 2020 (Figure 4C). This may have reduced the power in the GWAS due to missing flights during the 2018 juvenile vegetative growth stage, which reduced potential genetic variation, and unbalanced replications in 2020 due to weed pressure. However the range of MTA effect sizes increased in 2020 and showed the same amount of variation captured as in 2019 and 2021, which had more MTA (Table 2, Figure 4C). The conserved range of effect sizes without overlap between MTA indicates there is a great deal of variation in the genetic architecture of canopy cover in different growing conditions. Therefore, unique intra-year BLUPs for desired target environments may provide opportunities to overcome low heritability in certain years and identify previously undetected regions of the genome associated with a trait.

Distinct SNPs found by each modeling approach identified regions of the genome associated with specific environments or conditional GXE effects. As extreme weather events become more common and inter and intra-season variability increases, identifying germplasm capable of both maintaining performance and optimizing growth during favorable conditions will become essential to maintain yield and ensure global food production (Brown et al., 2015; Arias et al., 2021). By linking trait performance and marker effect size with environmental data using the types of modeling approaches highlighted in this study, researchers can generate an atlas of SNPs optimized for studying plant growth and stress-related functions (Della Coletta et al., 2023).

#### The interaction between environment and growth stage impacted regions of the genome associated with canopy cover

High temporal resolution data allowed the examination of how regions of the genome controlling maize canopy coverage changed throughout growth and development (Figure 4F). At a single GDD or GDD interval, there was an average of 33 MTA across all traits and modeling approaches. The GDDs with the most shared SNPs between percent and rate were 600 and 650, which also corresponded to the GDDs with the highest correlation between the two traits (Supplemental Figure S5). The range of effect sizes of unique SNPs found in each growth stage was constant, which demonstrates that there were equivalent amounts of genetic variation captured (Figure 4E). The temporal diversity of MTA observed for maize canopy cover matched the genetic architecture found by similar studies. Across 12 UAV flights, Rodene et al. (2022) identified nine unique regions of the maize genome with more than five significant SNPs in a 100 kb region involved in hybrid canopy coverage.

The different modeling approaches applied impacted the number of genomic regions detected at any given time (Figure 5A, 5B). Models that incorporated GxE estimates were successful at identifying SNPs in all growth stages, however there were GDD intervals with no significant associations. In 2019 and 2021, SNPs were identified using intra-year BLUPs at nearly all GDD, with a high MTA count at certain trait-time iterations, such as percent canopy cover in 2019 at 1100 and 1150 GDD. In contrast, intra-year models for 2018 and 2020 only detected SNPs in narrow intervals. These results show that regions of the genome involved with canopy coverage are highly dependent on the interaction between environment and developmental stage, which would help explain the sporadic distribution of MTA at discrete GDD and growth stages. These difference in SNPs detected by each model across time could also be due to differences in heritability and environmental discernibility in different GDD, which can cause BLUPs to shrink differently and reduce the ability to detect meaningful associations in individual environments compared to modeling approaches that incorporate data from multiple years (Van Tassel et al., 2022).

**Figure 5.**
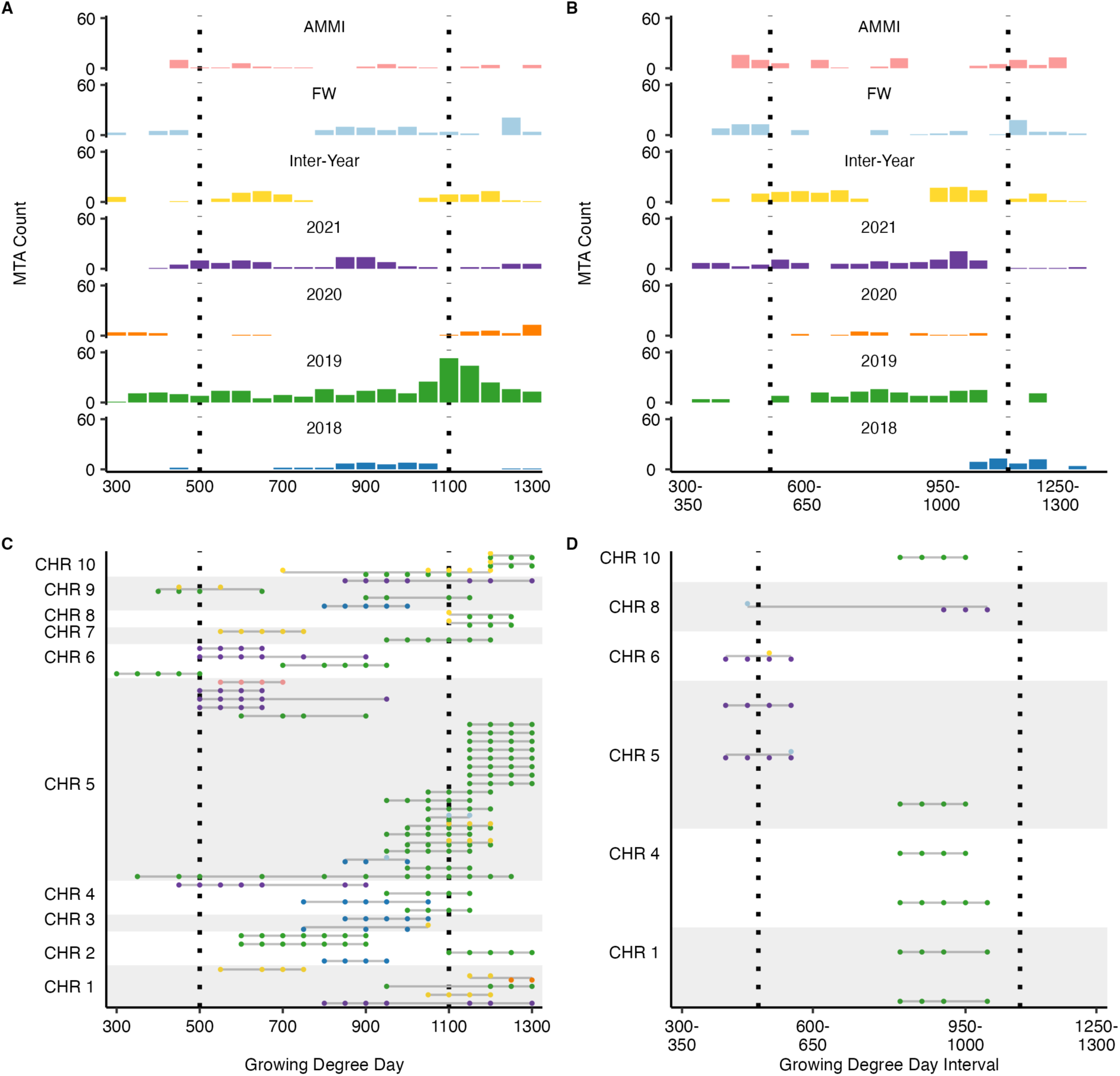
Temporal distribution and durability of single nucleotide polymorphisms (SNPs) associated with maize canopy coverage. A. The number of percent canopy cover marker trait associations (MTA) detected by each model [intra-year linear (2018, 2019, 2020, 2021), inter-year linear, Finlay-Wilkinson (FW) joint regression, and additive main-effects and multiplicative interaction (AMMI)] throughout the growing season. B. The rate of canopy development MTA detected by each model throughout the growing season. C. The temporally durable MTA with more than 3 occurrences across all models at separate growing degree days for percent canopy cover. The chromosome spacing is representative of the number of unique SNPs, not physical location. Gray bars are used to distinguish chromosomes. D. The temporally durable MTA with more than 3 occurrences at separate growing degree day intervals for canopy rate. Chromosome spacing is representative of the number of unique SNPs, not physical location. Gray bars are used to distinguish chromosomes.

#### Temporal GWAS was required to fully capture the genetic architecture of maize canopy cover

Due to complex interactions between genetics, phenology, and the environment, it can be difficult to reliably detect temporally durable regions of the genome involved with quantitative traits. Filtering GWAS results for temporally repeated SNPs results has been proposed as a useful method for reducing false discovery rates (Miao et al., 2019). In maize and soybean canopy cover and rice tiller angle, only one significant region for each trait was detected at multiple time points from temporal data sets (Xavier et al., 2017; Wu et al., 2019; Rodene et al., 2022). To identify temporally durable regions of the genome involved with maize canopy cover, we filtered for significant percent and rate of canopy cover SNPs with greater than three MTA across all modeling approaches and GDD (Knoch et al., 2020). Each separate modeling approach had at least one temporally durable SNP and in total there were 54 temporally durable significant SNPs for percent canopy cover and 10 for rate of canopy development (Figure 5C, 5D). Due to the repeatable significance and association with distinct environments or stability metrics, these temporally durable QTL would represent good candidates for gene validation efforts and further functional and physiological study.

There were less temporally durable regions of the genome for rate of canopy development (Figure 5D). This makes sense given the lower heritability and higher PVE by the environment at all growth stages for rate compared to percent canopy cover (Figure 2). It may be that SNPs associated with canopy growth rates simply change over time, resulting in less temporally durable SNPs. Alternatively, we observed oscillating variation in growth rates at a narrow time scale, which may indicate fluctuations between optimal and stressed growing conditions that could mask positive or negative genomic associations based on the environment at a given time. This could reduce the ability to reliably detect the same SNPs across multiple GDD intervals, despite substantial trait variation.

The temporal diversity of genomic regions involved in maize canopy coverage highlights the importance of time series data when studying quantitative, developmental traits. If percent canopy cover measurements had been restricted to a single, terminal time point at 1300 GDD, only 42 out of 393 unique significant SNPs and 15 out of 54 unique temporally durable significant SNPs would have been identified. From a breeding perspective, the temporal diversity of regions of the genome associated with canopy cover could allow favorable QTL to be stacked across growth stages to provide season-long benefits, such as increased weed suppression during the juvenile vegetative growth stage (Jannink et al., 2001). Temporal data and relevant genomic regions can also be used to dissect the functional mechanisms that contribute to canopy cover throughout development and compare putative genes that appear in narrow or wide time intervals.

## Conclusions

To understand the temporal phenomics of maize canopy cover, UAV images of the Wisconsin Diversity Panel throughout the growing season were collected and used to extract canopy cover percentage and growth rates using a k-means clustering algorithm and local polynomial regression modeling. When partitioning phenotypic variance across time, the range of primary and terminal maize canopy coverage was largely influenced by genetics, however, environmental conditions during the adult vegetative growth phase had a large effect on individual canopy accumulation. There was also a consistent impact of GxE interactions with different magnitudes and directions throughout development. Variation in canopy growth rates was influenced more by the environment than canopy percentage variation. Future research may choose to grow trials in more distinct environments and time UAV flights around potential causative weather events to further understand the intersection between environment and development on maize canopy cover.

When studying the genetic architecture of canopy cover, unique SNPs were found for distinct traits, modeling approaches, and time points. No single GDD, model, or trait captured the full genetic architecture contributing to diversity in maize canopy cover. More temporally durable regions of the genome were detected for SNPs associated with canopy percentage than rate, which may be due to the increased effect of intra-season variability on growth rates masking SNP effects across multiple GDDs. High throughput phenotyping is making the generation of multi-observational, big data sources possible in maize and other species of economic, health, and research significance. Continuing to increase the plant phenome with new biological, environmental, and temporal data is critical for understanding crop growth and development.

## Materials and Methods

### Experimental Design

In the 2018, 2019, 2020, and 2021 growing seasons, 501 selected maize lines from the Wisconsin Diversity Panel were planted at the University of Minnesota Agricultural Experiment Station located in St Paul, MN. Plants were grown in 15.5 foot single row plots, with 4 foot alleys and 30 in row spacing. Plots were seeded with a target of 27 seeds, for a density of 70,000 plants per hectare. Standard agronomic practices were employed. Planting dates were May 14, 2018, May 30, 2019, May 7, 2020, and May 6, 2021. Two replications of a randomized incomplete-block were planted each year, with replications divided into two blocks based on early (71-80 days after planting) and late (81-87 days after planting) flowering time. Inbred lines PH207 and B73 were included five times as check lines in each block. In total, 1040 plots were grown each year.

### UAV Image Acquisition and Image Processing

Aerial images of this experiment were collected at or near solar noon to minimize shadows using a DJI Phantom 4 Advanced in 2018 and 2019 and a DJI Phantom 4 RTK in 2020 and 2021. Flights were conducted at 30 m for a ground sampling distance of .82 cm with 80% front and side overlap to maximize reconstruction efficiency. Ground control points of known height and width were placed around the exterior of the field.

Between seven and twelve control points were installed each year, determined by field shape and neighboring plants. Between ten and twenty-seven image acquisition flights were conducted for each environment spanning May through September. Flight images were processed as previously described (Tirado et al., 2020). Briefly, crop service models and RGB colored orthomosaics were generated for each flight date using Agisoft Metashape Professional v1.7.5 software (Agisoft, 2022) (Figure 1A). A plot boundary raster layer was overlaid on each field level orthomosaic using QGIS v3.16 (QGIS Development Team, 2022). The 2019 environment suffered a severe lodging event, which disrupted plot delineations. This resulted in loss of observations during the middle of the growing season. To compensate for physical differences in plant locations as they recovered from the lodging event, a shifted plot boundary layer was used for the remainder of the 2019 flights.

### Canopy Cover Data Processing

To isolate plant objects within each plot, an unsupervised k-means clustering algorithm was implemented in MATLAB and Statistics Toolbox Release 2012b (The MathWorks, Inc.). This methodology clustered pixels within an image based on the similarity of their raw RGB values (Figure 1B). Each cluster was then manually classified as plant or non-plant. Clusters corresponding to soil, shadows, or other background field noise were grouped and removed from subsequent analysis (Figure 1C). Clusters classified as plant were masked within each orthomosaic (Figure 1D). Following masking, a dilation step was implemented in Matlab to add contiguity to images that had lost small plant pixels grouped into masked clusters (Figure 1E). A three pixel square dilation followed by a three or six pixel octagonal dilation was performed on each image using the ‘imclosè function in Matlab (Figure 1F). Size of the octagon dilation was dependent on the amount of missing plant pixels. This image processing pipeline was run on an Apple M1 Macbook Pro with 16GB of RAM.

### Data Quality Checks

For each plot, canopy cover was quantified as the percentage of masked plant pixels divided by the total number of pixels in each plot, as designated by the raster layer metadata. Weed pressure, stand count, and outlier canopy cover values were examined to quality check the data. Data quality control, analysis, and visualization was performed in R Studio Version 1.4.1717 using the ggplot2 and tidyverse packages (Wickham, 2016; Wickham et al., 2019; RStudio Team, 2020; R Core Team, 2022). The 2018 and 2020 growing season experienced significant weed pressure, and the k-means clustering could not distinguish between maize and weed plant tissue. Flight orthomosaic images were visually screened for weed pressure and plots with volunteer plants in their boundary were removed from the data set. This resulted in a loss of 114 plots in 2018 and 508 plots in 2020. No plots were removed due to weed pressure from 2019 or 2021. Stand counts were manually measured and ranged from 0-32 plants. A linear model was fit using the R lme4 package to test the effect of stand count on canopy cover (Bates et al., 2015). It was determined that a minimum of 13 plants was required to ensure statistically similar terminal canopy cover between plots. Plots with less than this minimum threshold were excluded from the analyses. Finally, outlier data points were identified as those with a deviation of 10% plant ratio deviations from the preceding and proceeding canopy cover values in a plot level growth curve. Individual data points considered outliers were removed from the growth curve, and a plot with more than three outliers was removed from subsequent analysis.

#### Growing Degree Day Calculations

To standardize maize growth phases between years, calendar dates were converted to GDD. Daily minimum and maximum temperatures and precipitation totals were recorded by University of Minnesota St. Paul weather station (Station ID: 218450). Daily GDD accumulation was calculated as ([*Temperature*_*max*_ − *Temperature*_*min*_] − 50)/2. Minimum and maximum temperatures less than 50°F and/or greater than 86°F were capped at 50°F and 86°F because maize growth is constrained beyond these thresholds.

### Canopy Cover Trait Prediction Using LOESS Regression

Following data cleaning protocols, a LOESS local polynomial regression was fit using the ‘loess’ function in the R ‘stats’ package to predict canopy coverage throughout the growing season (R Core Team, 2022). Spans from 0.15 to 0.95 were tested with 100 fold cross validation to determine the best fit for each environment based on the number of observations. A LOESS span of 0.25, 0.5, 0.3, and 0.3 was used for 2018, 2019, 2020, and 2021 respectively. For each environment, the smallest span while minimizing the mean error was chosen to preserve in-season variability for each growing condition. Variation in span depended on the number of flights and amount of missing data. From the LOESS regression curves, canopy cover was predicted from 300-1300 at 50 GDD intervals for each plot. In 2018 the predicted growth curve was shortened to 450-1300 GDD due to missing early season flights because LOESS curves cannot extrapolate beyond observed data points. Rate of canopy growth between each GDD interval was calculated as *b* = *y*_2_ − *y*_1_/*x*_2_ − *x*_1_, where y equals GDD and x represents canopy cover values.

### Analysis of Variance

For each trait and time combination, a multiple linear regression model fit using the ‘lm’ function of the R ‘lme4’ package was used to conduct ANOVA and determine significant experimental factors influencing canopy cover percentage and growth rate (Bates et al., 2015; R Core Team, 2022). For modeling PVE, the model *y*_*ijkl*_ = *μ* + *g*_*i*_ + *e*_*j*_ + *r*_(*j*)*k*_ + *b*_(*jk*)*l*_ + *ge*_*ij*_ + *ɛ*_*ijkl*_ was fit where *y*_*ijkl*_ is canopy cover or rate of canopy growth at a time, *μ* is the grand mean, *g*_*i*_ is the effect of genotype, *e*_*j*_ is the effect of year, *r*_(*j*)*k*_ is the effect of the *k*^*t*ℎ^ replication nested in the *j*^*t*ℎ^ environment, *b*_(*jk*)*l*_ is the effect of the *l*^*t*ℎ^ block nested in the *k*^*t*ℎ^ replication and *j*^*t*ℎ^ environment, *ge*_*ij*_ is the interaction between inbred and environment, and *ɛ*_*ijkl*_ is the residual. The PVE was calculated as the sum of squares divided by the sum of squares total from the ANOVA results. Heritability on a line mean basis for each GDD was calculated using the equation 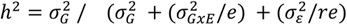 where 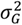 is the genotypic variance, 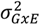 is the genotype by environment interaction variance, 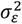 is the error variance, *e* is the number of environments, and *r* is the number of replications.

### Static and Dynamic Stability Estimations

Dynamic and static stability estimates were calculated for each canopy cover trait-time iteration. Low number of replications for a genotype can skew stability estimates and confound analysis. Therefore, stability estimates were only obtained for genotypes with a minimum of five replications grown over a minimum of three environments. The “ammistability” package in R Studio Version 1.4.1717 was used to calculate static stability metrics using an AMMI model (Aravind and Ajay, 2018). All significant principal components for each phenotype and time point were used in the calculations, however in the absence of multiple significant principal components a minimum of two was enforced for use in the AMMI analysis. AVAMGE quantifies static stability as the sum of significant principal components and associated eigenvalues. The “FW” package in R Studio Version 1.4.1717 was utilized to quantify dynamic stability (Lian and de Los Campos, 2015). A total of 50,000 iterations of a Gibbs sampling algorithm was run for each FW regression, with 5,000 burn-in to allow distributions to normalize.

### Best Linear Unbiased Predictions of Canopy Cover Traits

For percent and rate traits, BLUPs were calculated at each time point using the ‘lm’ function of the R ‘lme4’ package (Bates et al., 2015; R Core Team, 2022). Both intra and inter year BLUPs were calculated. To avoid overfitting and preserve inter-environment variation, if replication or block was not significant (p < .05) for a trait-time iteration, the factor was removed from the model prior to BLUP calculation. The complete intra year BLUP model was *y*_*ikl*_ = *μ* + *g*_*i*_ + *r*_*k*_ + *b*_(*k*)*l*_ + *ɛ*_*ikl*_ where *y*_*ikl*_ is a given trait-time combination, *μ* is the grand mean, *g*_*i*_ is the effect of genotype, *r*_*k*_ is the effect of the *k*^*t*ℎ^ replication, *b*_(*k*)*l*_ is the effect of the *l*^*t*ℎ^ block nested in the *k*^*t*ℎ^ replication, and *ɛ*_*ikl*_ is the residual. When replication was not significant, this model was simplified to *y*_*il*_ = *μ* + *g*_*i*_ + *b*_*l*_ + *ɛ*_*il*_ where *b*_*l*_ is the lth block, not nested in replication. If block was not significant, the model was simplified to *y*_*ik*_ = *μ* + *g*_*i*_ + *r*_*k*_ + *ɛ*_*ik*_. If neither block nor replication was significant, the model was simplified to *y*_*i*_ = *μ* + *g*_*i*_ + *ɛ*_*i*_. The inter-year BLUPs were calculated using the same model used in ANOVA for PVE. When replication was not significant, this model was simplified to *y*_*ijkl*_ = *μ* + *g*_*i*_ + *e*_*j*_+ *b*_(*j*)*l*_ + *ge*_*ij*_ + *ɛ*_*ijl*_. When block was not significant, the model was simplified to *y*_*ijk*_ = *μ* + *g*_*i*_ + *e*_*j*_ + *r*_(*j*)*k*_ + *ge*_*ij*_ + *ɛ*_*ijk*_. If neither block nor replication was significant, the model was simplified to *y*_*ij*_ = *μ* + *g*_*i*_ + *e*_*j*_ + *ge*_*ij*_ + *ɛ*_*ij*_.

### Genome-Wide Association Analysis

A GWAS was performed to find MTA for all trait-model-time phenotypes as previously described (Burns et al., 2021; Renk et al., 2021). Intra- and inter-year BLUPs, AVAMGE values, and FW slopes were used as phenotypes. Marker values from Qui et al. (2021) and filtered by Renk et al. (2021) were used as genotypic data. Briefly, GAPIT v.3 was used to convert genomic data into numeric format and generate a genetic map and PCA covariate values (Wang and Zhang, 2021). The first five PCAs were used. Alleles with a minor allele frequency less than 5% were excluded from the GWAS. The Fixed and random model Circulating Probability Unification (FarmCPU) method was implemented in GAPIT v.3 for each trait-year/stability-time BLUP dataset (Liu et al., 2016; Wang and Zhang, 2021). The genetic map and filtered numeric genomic and BLUP datasets were used to generate a suggested p-value for marker significance based on 100 permutations of the FarmCPU model. A linkage disequilibrium matrix for the genomic data was calculated using PLINK v.1.90b6.10 (Purcell et al., 2007). Linkage disequilibrium blocks of 1 MB were created, excluding markers with >25% missing data. A minimum threshold of .9 was then used to merge significant SNPs in linkage disequilibrium. Effects sizes were normalized across traits, models, and time by dividing raw marker effects by the range of phenotypic values used in the corresponding trait-model-time GWAS. SNP location and nearby genomic elements are based on the B73 v5 reference genome assembly and annotation (Hufford et al., 2021; Woodhouse et al., 2021).

## Acknowledgments

We thank the Minnesota Supercomputing Institute at the University of Minnesota for providing resources that contributed to the research results reported in this article.

## Author contributions

Conceptualization: C.N.H and C.D.H. Data curation: J.C., D.D.S., S.B.T. Formal analysis: J.C. Funding acquisition: J.C., D.D.S., S.B.T., N.M.S., C.N.H., C.D.H. Methodology: J.C., C.N.H, C.D.H., Software: J.C., D.D.S., S.B.T. Writing - original draft: J.C., C.N.H, C.D.H. Writing - review and editing: J.C., D.D.S., N.M.S., C.N.H., C.D.H.

## Supplemental data

**Supplemental Table S1.** Percent canopy cover values at each local regression predicted time point from 300-1300 GDD for each genotype in each year.

**Supplemental Table S2.** Mean, minimum, maximum, standard deviation (SD) and coefficient of variation (CV) of percent canopy cover at 50 GDD intervals, rate of canopy development between 50 GDD intervals, and for each trait averaged across growth stages. Percent of variance explained (PVE) by factors influencing percent canopy cover at 50 GDD intervals, rate of canopy development between 50 GDD intervals, and PVE for each trait averaged across growth stages. Insignificant factors (p > .05) are asterisked. Mean, minimum, maximum, standard deviation (SD) and coefficient of variation (CV) of the sum across environments of absolute value of GxE interaction (AVAMGE) modeled by an additive main-effects and multiplicative interaction (AMMI) model for genotypes using percent canopy cover at 50 GDD intervals, rate of canopy development between 50 GDD intervals, and each trait averaged across growth stages. Mean, minimum, maximum, standard deviation (SD) and coefficient of variation (CV) of Finley-Wilkinson (FW) regression coefficients for genotypes using percent canopy cover at 50 GDD intervals, rate of canopy development between 50 GDD intervals, and each trait averaged across growth stages

**Supplemental Table S3.** Significant single nucleotide polymorphisms (SNP), chromosome number and position, p-value, minor allele frequency (maf), normalized effect size between phenotype-model-GDD iterations, phenotype [percent canopy cover (Percent) and rate of canopy development (Rate)], model [intra-year linear (2018, 2019, 2020, 2021), inter-year linear, Finlay-Wilkinson (FW) joint regression, and additive main-effects and multiplicative interaction (AMMI)], predicted growing degree day (GDD) or GDD interval, and growth stage [juvenile vegetative (Juv.), adult vegetative (Adult), and reproductive (Rep.)]

**Supplemental Figure S1.** Observed and predicted canopy cover for randomly sampled B73 plots from each environment. Each X represents the canopy cover at the growing degree day of a UAV flight. Circles indicate local regression predicted canopy cover values at 50 growing degree day intervals. Juvenile vegetative, adult vegetative, and reproductive growth stages are separated by dashed black lines.

**Supplemental Figure S2.** Weather trends and percent and rate of canopy coverage in each environment. Juvenile vegetative, adult vegetative, and reproductive growth stages are separated by dashed black lines. A. Seasonal temperature and precipitation in each environment. Minimum daily temperatures are shown in light blue and maximum daily temperatures are shown in red. Precipitation events greater than 0.1 inches are shown in blue. B. Distribution of plot percent canopy coverage at local regression predicted 50 growing degree day intervals. A smoothed curve was fit for each environment. C. Distribution of plot rate of canopy development between local regression predicted 50 growing degree day intervals. A smoothed curve was fit for each environment.

**Supplemental Figure S3.** Growth curves of representative genotypes with high and low static and dynamic stability. Juvenile vegetative, adult vegetative, and reproductive growth stages are separated by dashed black lines from left to right. A. Representative inbred I29 with low static stability modeled by an additive main-effects and multiplicative interaction (AMMI) model across all Growing Degree Days. B. Representative inbred PHM10 with high static stability modeled by an AMMI model across all Growing Degree Days. C. Representative inbred YE 4 showing low dynamic stability based on Finlay-Wilkinson (FW) slope coefficients across all Growing Degree Days. D. Representative inbred N192 showing high dynamic stability based on FW slope coefficients across all Growing Degree Days.

**Supplemental Figure S4.** Correlation between models taking into consideration genotype by environment interactions for canopy cover. Juvenile vegetative, adult vegetative, and reproductive growth stages are separated by dashed black lines. A. Correlation between inter-year best linear unbiased predictions, Finlay-Wilkinson (FW) slope coefficients, and additive main-effects and multiplicative interaction (AMMI) model values for percent canopy cover. B. Correlation between inter-year BLUPs, FW slope coefficients, and AMMI values for rate of canopy development.

**Supplemental Figure S5.** Correlation between percent canopy cover at 50 growing degree day intervals and rate canopy cover development between 50 growing degree day intervals combined from all environments. Juvenile vegetative, adult vegetative, and reproductive growth stages are separated by dashed black lines.

**Supplemental Figure S6.** Count of marker trait associations (MTA) across all phenotypes, models, and growing degree days for each unique significant single nucleotide polymorphism (SNP).

## Data Availability

Scripts and files used to generate and analyze data are available on GitHub at https://github.umn.edu/cdhirschLab/Julian/tree/master/canopy_cover. All data including the UAV-derived canopy cover values, orthomosaics, DEMs, plot boundary files, and mask files for each date of UAV data collection, cumulative GDDs calculated for each date of data collection, and weather data has been made available at the Digital Repository for U of M (DRUM) at https://doi.org/10.13020/SKJN-QX31.

