## Supplemental Figure S1 for "Dissecting the temporal phenomics and genomics of maize canopy cover using UAV mediated image capture"

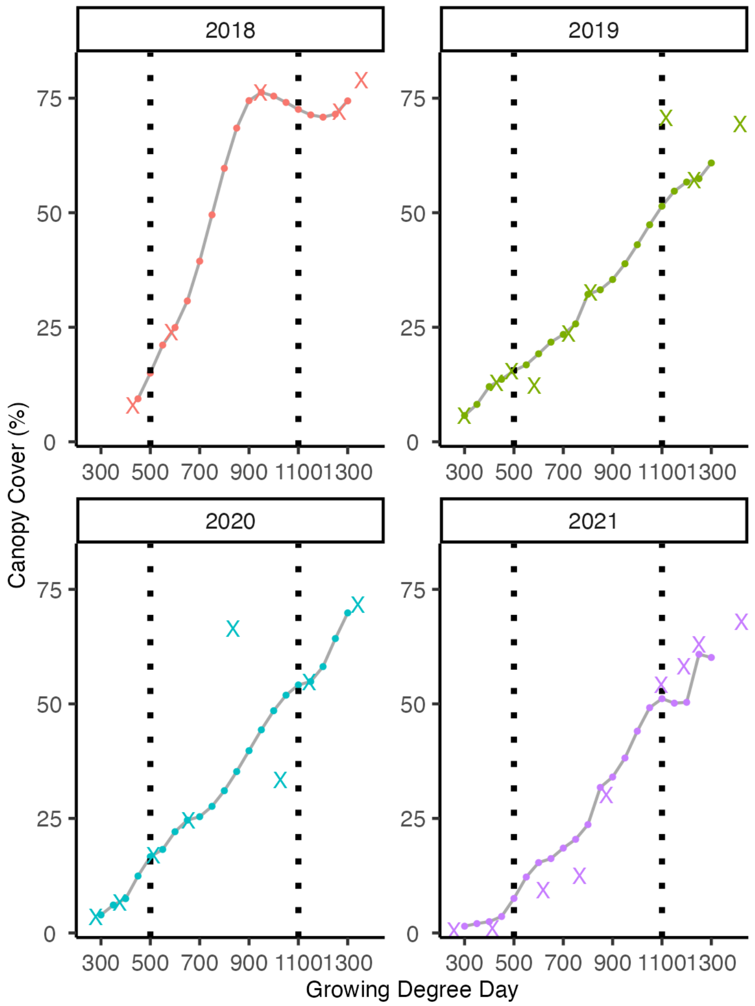


**Supplemental Figure S1.** Observed and predicted canopy cover for randomly sampled B73 plots from each environment. Each X represents the canopy cover at the growing degree day of a UAV flight. Circles indicate local regression predicted canopy cover values at 50 growing degree day intervals. Juvenile vegetative, adult vegetative, and reproductive growth stages are separated by dashed black lines.
