## Supplemental Figure S2 for "Dissecting the temporal phenomics and genomics of maize canopy cover using UAV mediated image capture"

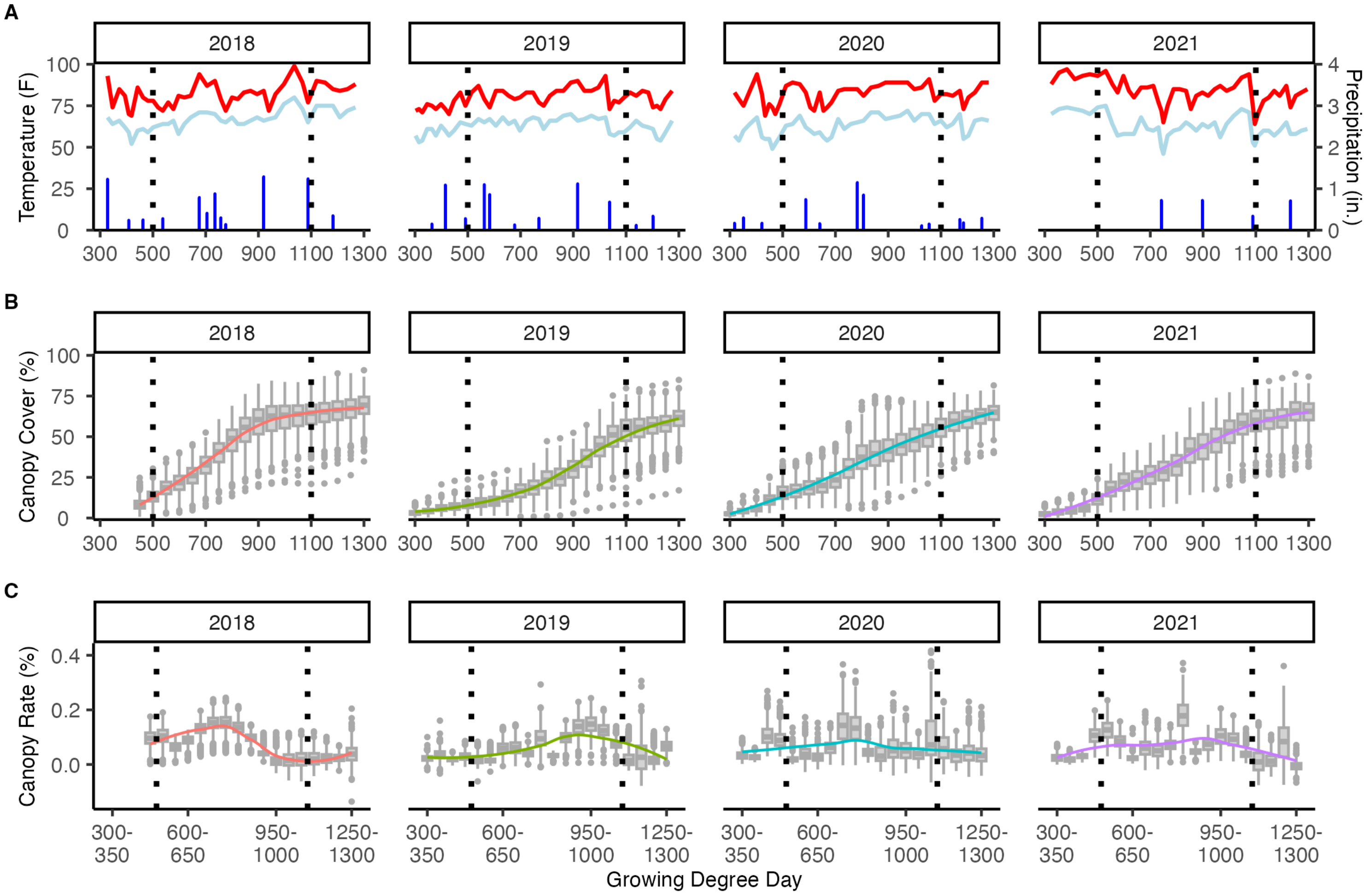


**Supplemental Figure S2.** Weather trends and percent and rate of canopy coverage in each environment. Juvenile vegetative, adult vegetative, and reproductive growth stages are separated by dashed black lines. A. Seasonal temperature and precipitation in each environment. Minimum daily temperatures are shown in light blue and maximum daily temperatures are shown in red. Precipitation events greater than 0.1 inches are shown in blue. B. Distribution of plot percent canopy coverage at local regression predicted 50 growing degree day intervals. A smoothed curve was fit for each environment. C. Distribution of plot rate of canopy development between local regression predicted 50 growing degree day intervals. A smoothed curve was fit for each environment.
