## Supplemental Figure S3 for "Dissecting the temporal phenomics and genomics of maize canopy cover using UAV mediated image capture"

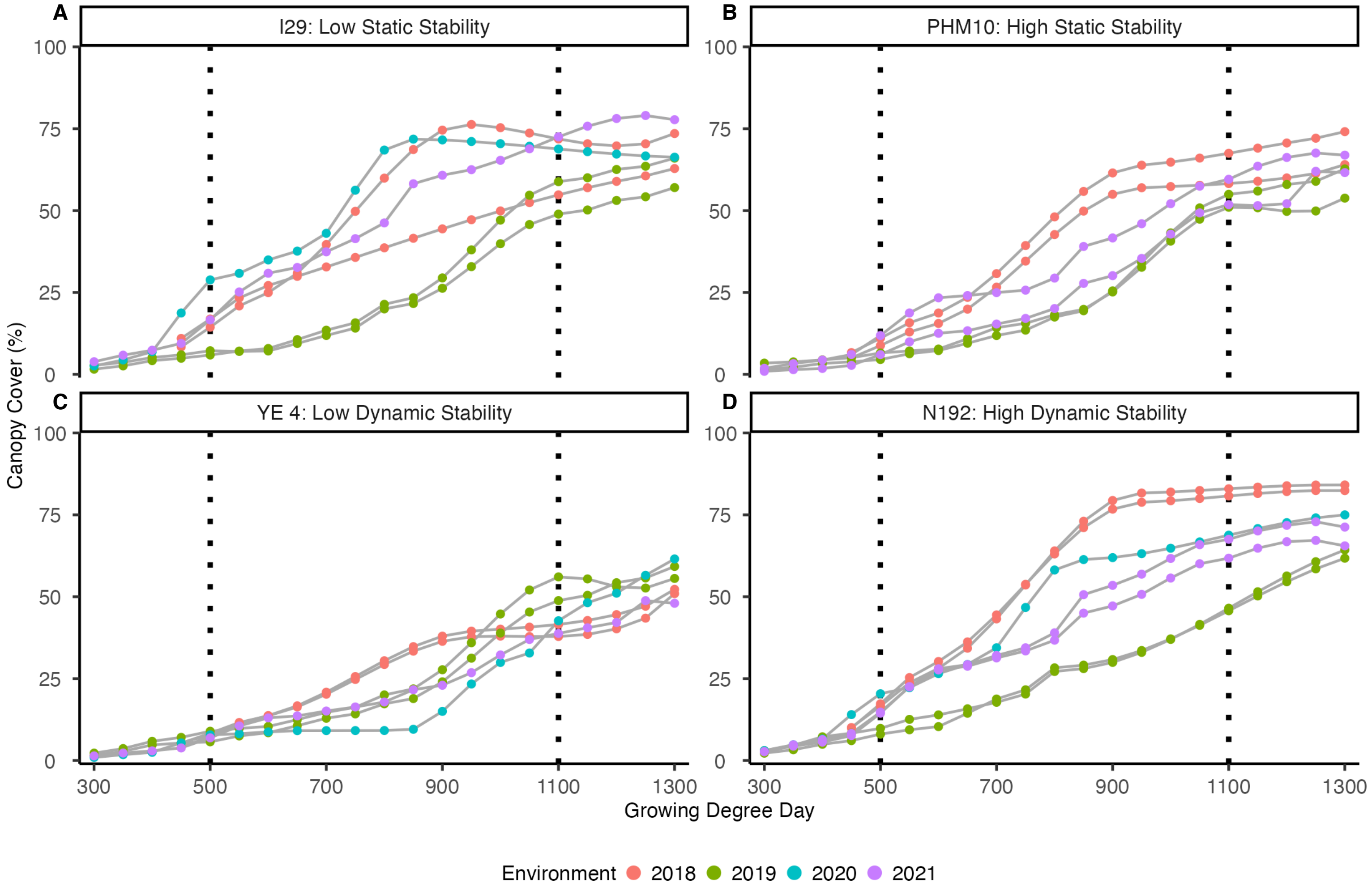


**Supplemental Figure S3.** Growth curves of representative genotypes with high and low static and dynamic stability. Juvenile vegetative, adult vegetative, and reproductive growth stages are separated by dashed black lines from left to right. A. Representative inbred I29 with low static stability modeled by an additive main-effects and multiplicative interaction (AMMI) model across all Growing Degree Days. B. Representative inbred PHM10 with high static stability modeled by an AMMI model across all Growing Degree Days. C. Representative inbred YE 4 showing low dynamic stability based on Finlay-Wilkinson (FW) slope coefficients across all Growing Degree Days. D. Representative inbred N192 showing high dynamic stability based on FW slope coefficients across all Growing Degree Days.
