## Supplemental Figure S4 for "Dissecting the temporal phenomics and genomics of maize canopy cover using UAV mediated image capture"

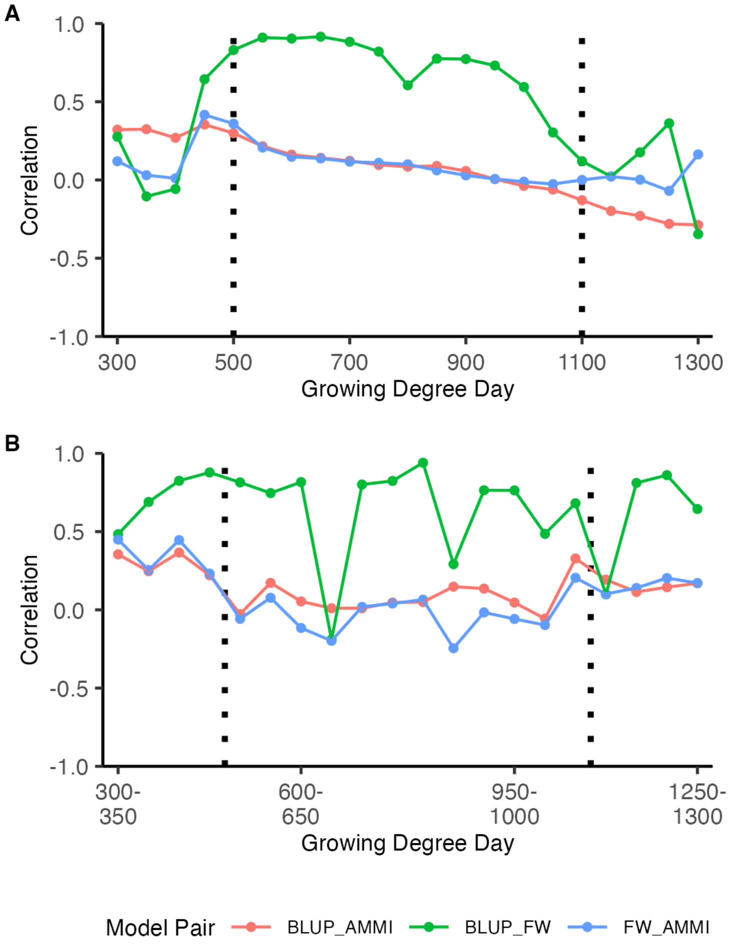


**Supplemental Figure S4.** Correlation between models taking into consideration genotype by environment interactions for canopy cover. Juvenile vegetative, adult vegetative, and reproductive growth stages are separated by dashed black lines. A. Correlation between inter-year best linear unbiased predictions, Finlay-Wilkinson (FW) slope coefficients, and additive main-effects and multiplicative interaction (AMMI) model values for percent canopy cover. B. Correlation between inter-year BLUPs, FW slope coefficients, and AMMI values for rate of canopy development.
