## Supplemental Figure S5 for "Dissecting the temporal phenomics and genomics of maize canopy cover using UAV mediated image capture"

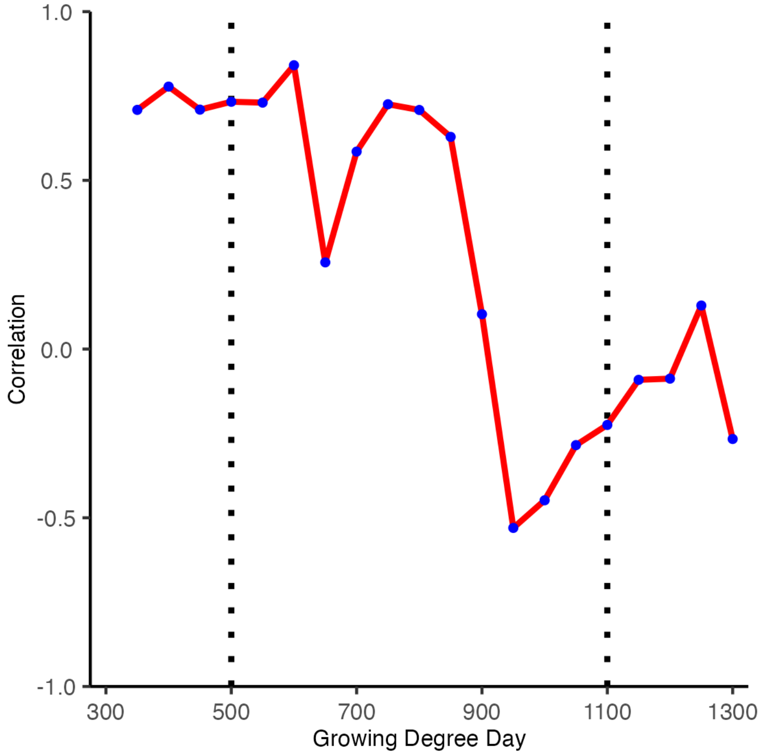


**Supplemental Figure S5.** Correlation between percent canopy cover at 50 growing degree day intervals and rate canopy cover development between 50 growing degree day intervals combined from all environments. Juvenile vegetative, adult vegetative, and reproductive growth stages are separated by dashed black lines.
