## Supplemental Figure S6 for "Dissecting the temporal phenomics and genomics of maize canopy cover using UAV mediated image capture"

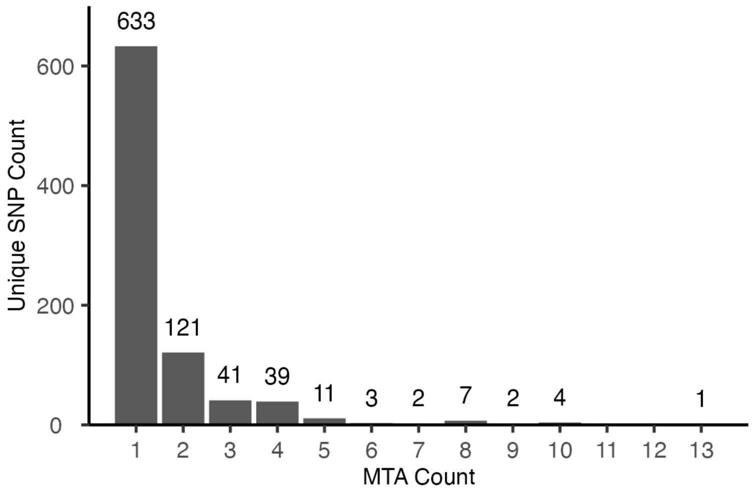


**Supplemental Figure S6.** Count of marker trait associations (MTA) across all phenotypes, models, and growing degree days for each unique significant single nucleotide polymorphism (SNP).
